# Fusion expression and anti-*Aspergillus flavus* activity of a novel inhibitory protein DN-AflR

**DOI:** 10.1101/302869

**Authors:** Yuan Liang, Qing Kong, Yao Yao, Shujing Xu, Xiang Xie

## Abstract

The regulatory gene (*aflR*) of aflatoxin encodes AflR, a positive regulator that activates transcriptional pathway of genes in aflatoxin biosynthesis. New L-Asp-L-Asn (DN) extracted from *Bacillus megaterium* inhibited the growth of *A. flavus* had been elucidated in our laboratory. The genes encoding DN and binuclear zinc finger cluster protein of AflR were fused, then fusion protein could compete with the AflS-AflR complex for the AflR binding site and significantly improve anti-*A. flavus* activity of DN. The fusion gene *dn-aflR* was cloned into pET32a and recombinant plasmid was introduced into *Escherichia coli* BL21. The highest expression was observed after 10 h induction and purified by affinity chromatography column. Compared with DN, the novel fusion protein DN-AflR significantly inhibited the growth of *A. flavus* and biosynthesis of aflatoxin B_1_. This study promoted the use of competitive inhibition of fusion proteins to reduce the expression of regulatory genes in the biosynthetic pathway of aflatoxin. Moreover, it provided more supports for deep research and industrialization of such novel, anti-*A. flavus* bio-inhibitors.

**IMPORTANCE:** Aflatoxin contamination has seriously influence on export of agricultural products, income of farmers and economic development. Biological methods, especially using antagonistic microorganisms to inhibit aflatoxin biosynthesis gradually become the hot spot in recent years. DN (L-Asp-L-Asn) from *Bacillus megaterium*, which could inhibit growth of *Aspergillus flavus* and synthesis of aflatoxin, has been identified. In this report, we fused the genes encoding inhibitory peptides (DN) and specific zinc finger cluster protein, and expressed the novel anti-*A*. *flavus* protein in *Escherichia coli*. Compared with DN, the inhibitory ability of novel protein has been improved significantly. This research showed fusion expression of anti-fungal proteins, such as DN-AflR, is a promising method to economically improve the inhibitory activity of bio-inhibitors for *A. flavus*.

## INTRODUCTION

Aflatoxin is one of the most potent naturally occurring toxic and carcinogenic compound, which is a mycotoxin that poses a serious threat to human health (1). Aflatoxin contamination has seriously influence on export of agricultural products, income of farmers and economic development (2, 3). Biological methods, especially using antagonistic microorganisms to inhibit aflatoxin biosynthesis gradually become hot spot in recent years (4). Palumbo et al. isolated one strain of *Bacillus* from almonds, which could inhibit the growth of *Aspergillus flavus* (5). Early studies found that *Mycobacterium smegmatis* and *Rhodococcus erythropolis* could produce F420 H2-dependent reductases to degrade aflatoxin (6, 7). Some studies isolated antifungal compounds from *Bacillus* and verified their inhibitory effects on the growth of *A. flavus* (8, 9). Above all, antagonistic microorganisms could produce metabolites or enzymes to inhibit expression of regulatory genes, or degrade aflatoxin.

Aflatoxin biosynthetic pathway has been studied for years and is one of the best understood fungal secondary metabolic pathways. The whole-genome sequencing of *A. flavus* has been accomplished, and we could better control aflatoxin contamination through deep research of the regulatory genes and mechanisms. Up to now, at least 34 genes have been identified as members of the aflatoxin pathway gene cluster. On the 82-kb biosynthetic gene cluster of aflatoxin, *aflR* and *aflS* (formerly known as *aflJ*) are genes involved in pathway regulation (10, 11). *aflR* is necessary for transcription of some genes in *Aspergillus* gene cluster (12–14). This gene encodes a specific DNA binding protein (AflR) containing 444 amino acids, and the 29th to 56th amino acids constitute a binuclear zinc finger cluster protein with sequence Cys-Xaa2-Cys-Xaa6-Cys-Xaa6-Cys-Xaa2-Cys-Xaa6-Cys, which is the key region of AflR to activate gene transcription and determines AflR-binding specificity (15). AflR, which possesses DNA-binding and activation domains typical of the GAL4-type family of positive regulatory proteins in yeast and fungi (16). A common feature of fungal gene clusters, including those for secondary metabolites, is the presence of specific regulatory genes, which have been found to encode members of the zinc binuclear cluster protein family typified by GAL4 (17–19). AflS was found to be involved in the regulation of transcription. Between *aflR* and *aflS* is intergenic region, where promoter region is located. AflS-AflR complex binds to AflR binding site and activates aflatoxin biosynthesis. Studies suggested that failure to product aflatoxin due possibly to alternation in the interaction between AflS and AflR (10, 20). Bio-inhibitors inhibited expression of *aflR* or *aflS* mainly by acting on the intergenic region, thereby prevented the formation of AflS-AflR complex, so that AflS-AflR complex could not bind to AflR binding site and activate biosynthetic pathway of aflatoxin.

Recently, we identified DN (L-Asp-L-Asn) from *Bacillus megaterium*, which could inhibit growth of *A. flavus* and synthesis of aflatoxin. To improve the inhibitory effect of DN, we transformed the genes encoding DN and GAL4-type zinc finger cluster protein (specifically binds to the AflR binding site) by gene fusion. We hypothesized the fusion protein could compete with AflS-AflR complex by acting on AflR binding sites to inhibit the activation of AflS-AflR complex and improve anti-*A. flavus* activity of DN.

## RESULTS

### Effects of DN on mycelia morphology of *A. flavus*. (i) TEM

In control group, the internal structure of cells grew normally and various organelles were clearly visible (Fig. 2A and B). However, cell structures of mycelia treated by DN were obviously abnormal: organelles were degenerated, and vacuole became significantly larger, expanded and fractured (Fig. 2C and D).

**FIG 1.**
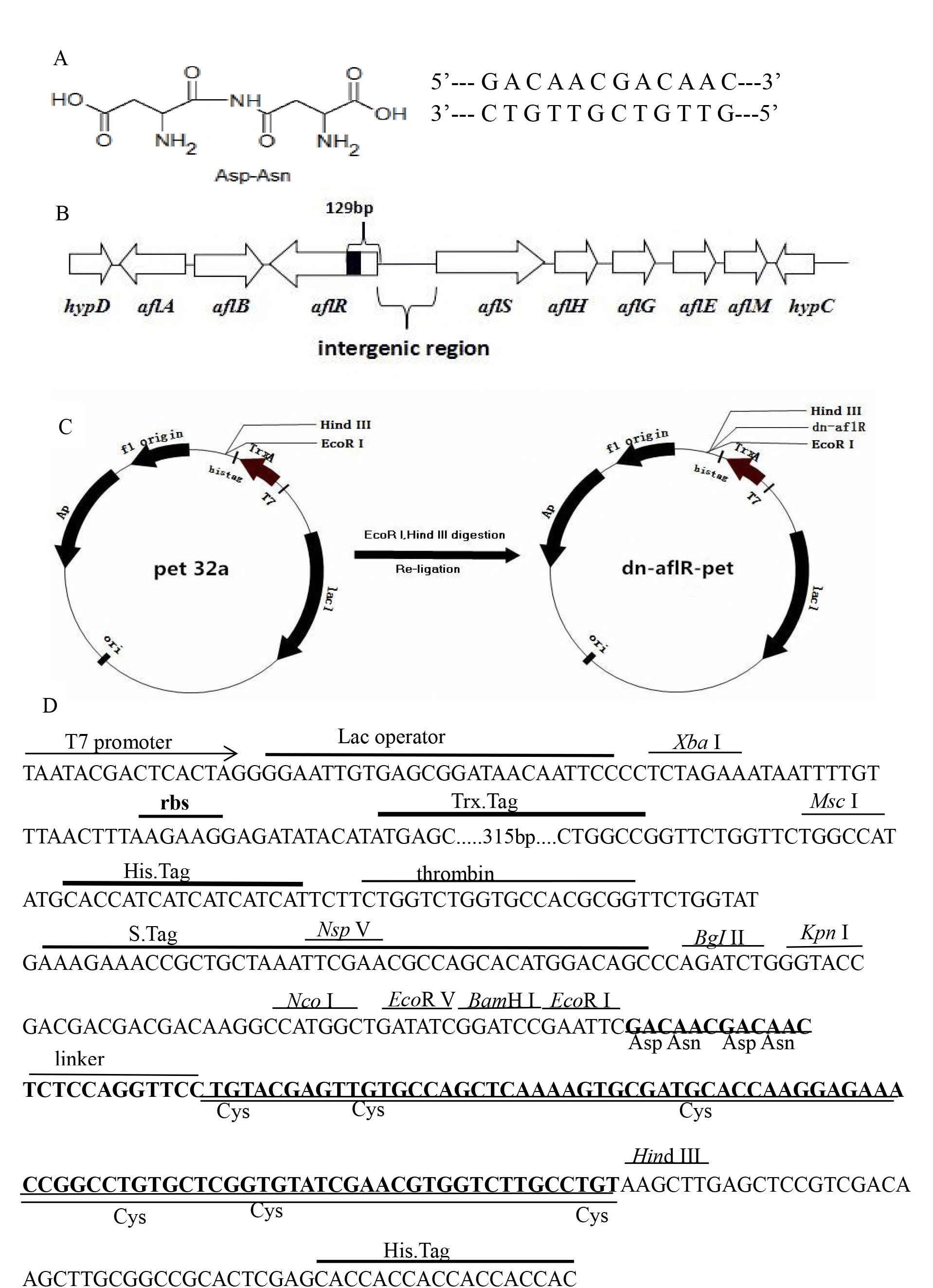
Molecular modified fragments and characteristics of *E. coli* expression plasmid dn-aflR-pet32a. (A) The chemical formula and base sequences of DN. (B) The aflatoxin biosynthetic pathway and clustered genes, AflR binding site (black area) and intergenic region. (C) Construction of expression plasmid. DNA fragment was digested with Hind III and EcoR I, and inserted into downstream of the thioredoxin (Trx) and then ligated with T4 DNA ligase. DN-AflR was expressed as a fusion protein with Trx and 6His. Ar, an ampicillin resistance gene. (D) Base composition of the inserted fusion gene (thick base) containing genes encoding DN (under line) and zinc finger cluster protein (double line). 6His tag, Trx tag and unique restriction sites (over line) were also indicated.

**FIG 2.**
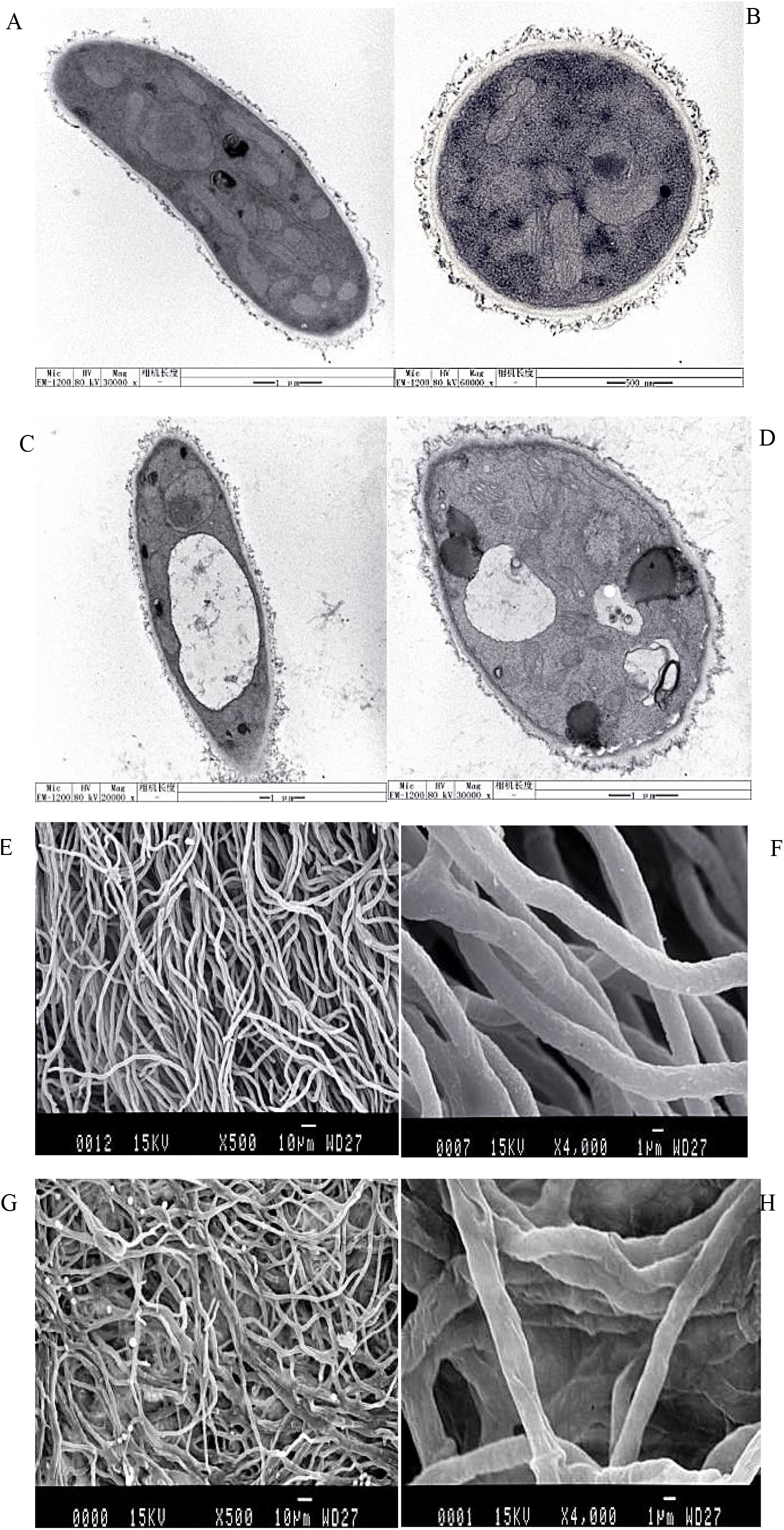
Effects of DN on mycelia morphology of *A. flavus*. (A), (B), (C), (D): TEM; (E), (F), (G), (H): SEM; (A), (B), (E), (F): Control groups; (C), (D), (G), (H): DN treatment groups.

### (ii) SEM

Mycelia of control group were integrity, showing straight, neatly arranged and smooth surface (Fig. 2E and F). While mycelia treated with DN grew abnormally: mycelia were rough and had uneven thickness, and appeared to be distorted and broken (Fig. 2G and H).

### Expression and identification of fusion protein

The expected DNA and vector fragments were seen in 1% agarose electrophoresis after digesting with EcoR I and Hind III. Colony PCR and DNA sequencing showed that target fragment (*dn-aflR*) was successful inserted into the vector. The molecular weight of empty plasmid (Trx-His-pET32a) and fusion protein (Trx-His-DN-AflR) were predicted to be 19 kDa and 22 kDa. Expression vector with fusion gene had a distinct protein band (24 kDa, Fig. 3A, lanes 1 and 2). The empty plasmid did not express the band at the corresponding position and had its own specific band (Fig. 3A, lane 3). Results illustrated successful expression of the fusion protein DN-AFLR. The supernatant after ultrasonic breaking (Fig. 3B, lane 1) and the precipitate after 8 mM urea dissolved (Fig. 3B, lane 2) both showed obvious protein bands. The result was consistent with predicted weight of expected protein before purification (Fig. 3A). There were no obvious protein bands in wash buffer (Fig. 3B, lanes 3-5), and there was only one obvious protein band in elution buffer (Fig. 3B, lanes 6 and 7), which corresponded to the target protein. ImageJ software (https://imagej.nih.gov/ij/) was used to compare the density of the bands on the gel, the fusion protein accounts for approximately 57.4% of the total cellular protein, the relative content of fusion protein was about 600 ug ml^−1^.

**FIG 3.**
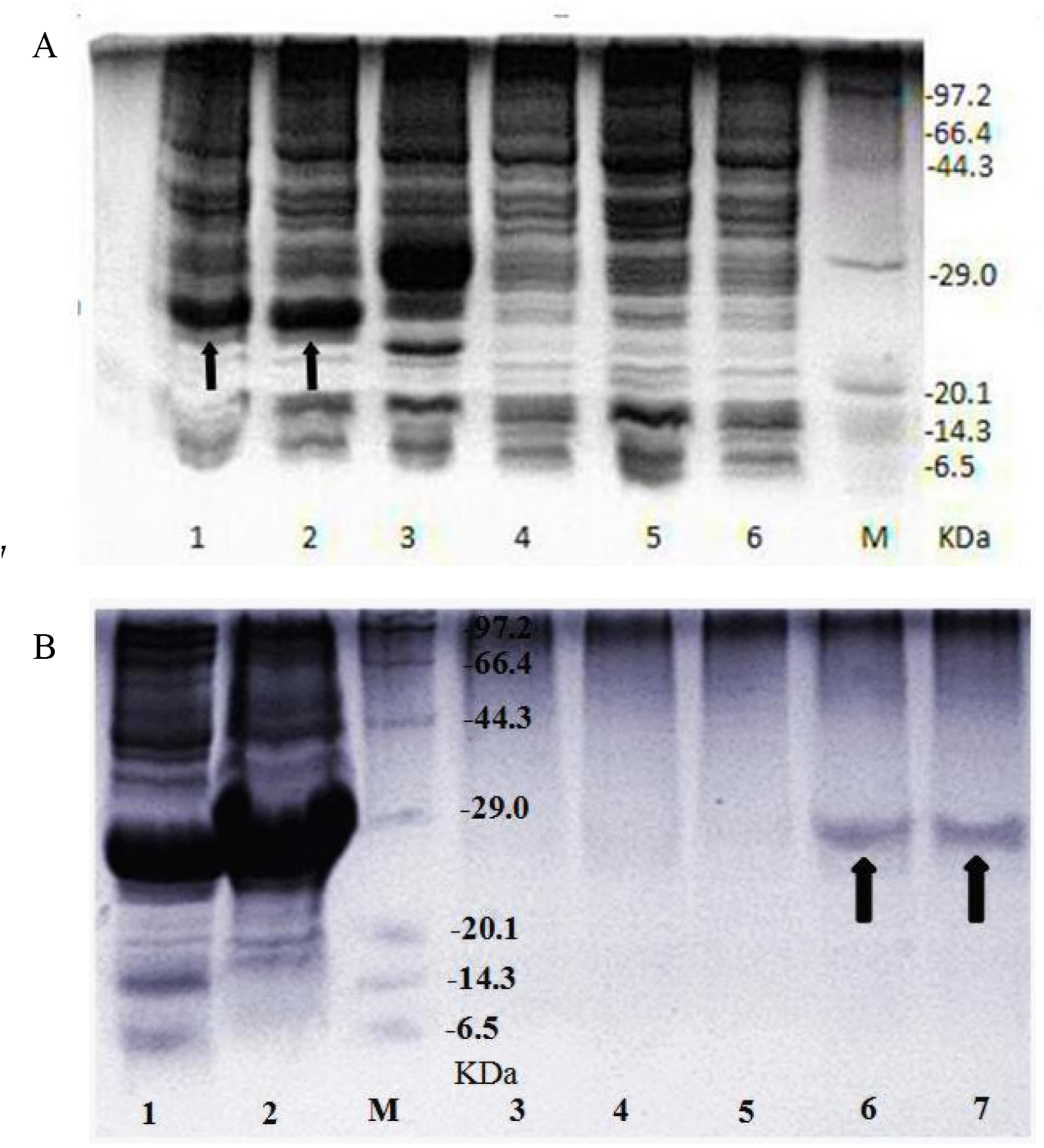
SDS-PAGE analyzed the expression of DN-AflR. (A) M: molecular weight marker (Takara). Lanes 1 and 2: protein bands of *dn-aflR-pET32a* expression. Lane 3: protein band of empty pET32a expression. Lanes 4, 5 and 6: negative control (before inducted). The position of the fusion protein was indicated with an arrowhead. (B) Analysis of purified fusion protein DN-AflR. M: marker. Lane 1: expressed supernatant after ultrasonic breaking. Lane 2: precipitate after 8mM urea dissolved. Lanes 3, 4 and 5: 50 mM imidazole buffer band, 100 mM imidazole buffer band and 300mM imidazole buffer band, respectively. Lanes 6 and 7: 200mM imidazole elution buffer band. The position of the purified protein was indicated with an arrowhead.

### Inhibitory effect of DN-AflR on growth of *A. flavus*

Plate treated by DN had only a small inhibition zone of 5.0±0.2 mm (Fig. 4A, plate a). The control group (Fig. 4A, plate c) and 200 mM imidazole treated group (Fig. 4A, plate b) showed no significant zone of inhibition. Diameter of inhibition zone treated with fusion protein was as high as 21.3±0.2 mm (Fig. 4A, plates d-f). Inhibitory ability of DN-AflR was much stronger than that of DN. The MICs of DN-AflR and DN were 150 μg ml^−1^ and 400 μg ml^−1^, respectively. MFC of DN-AflR was 400 μg ml^−1^. Results of two repeated experiments were consistent (Table 1). The MIC and MFC of fusion protein were lower than that of DN, indicating that the fusion protein had stronger inhibitory effect on growth of *A. flavus* at a lower concentration.

**FIG 4.**
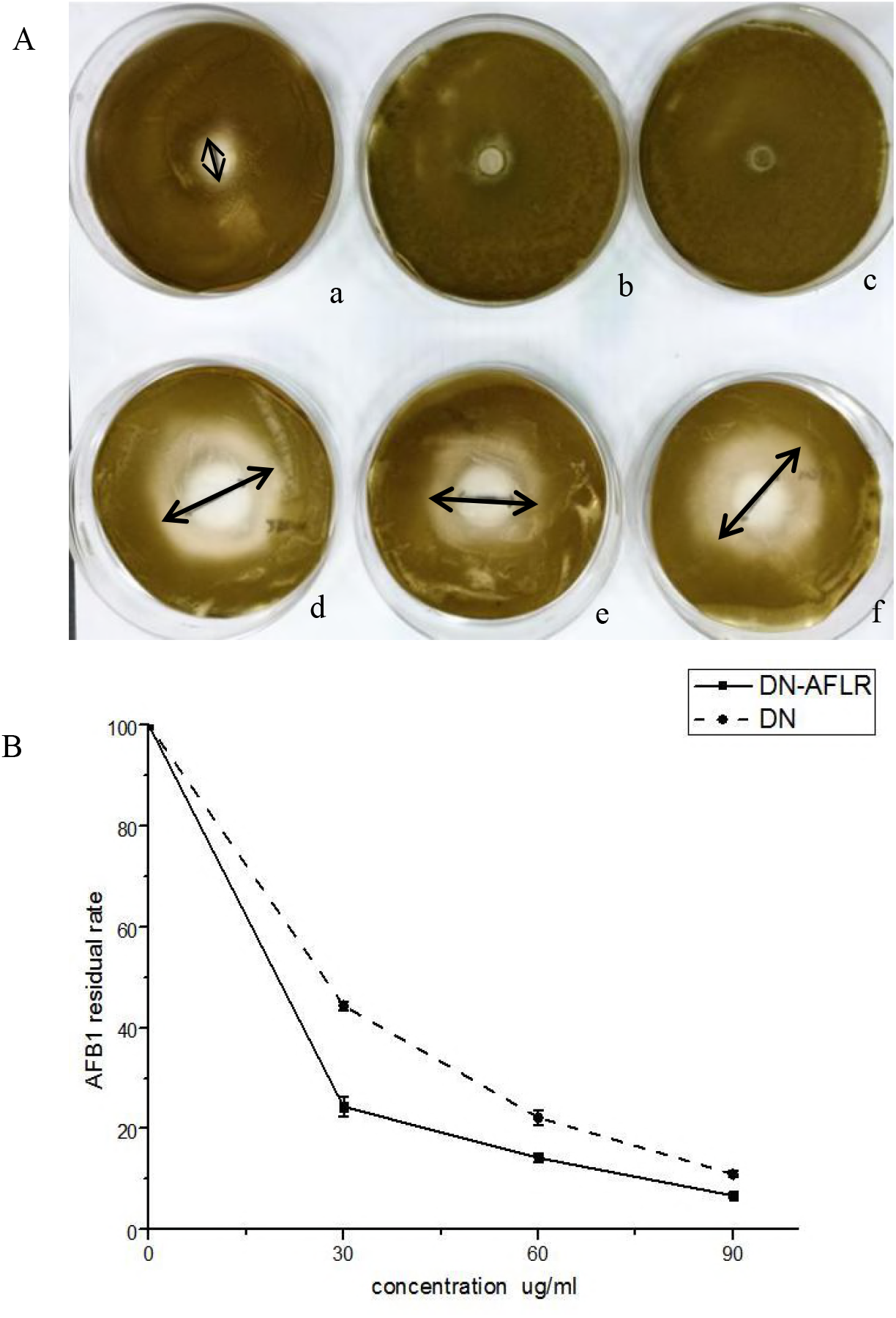
Anti-*A. flavus* activity and degradation of AFB_1_. (A) Plate a: treated with DN. Plate b: treated with 200 mM imidazole. Plate c: control group with no treatment. Plates d, e and f: treated with DN-AflR, the circle diameters of them were around 24, 19 and 21mm, respectively. (B) The abscissa represented the final concentration of DN (solid line) and DN-AflR (dotted line), the ordinate (content of AFB_1_ without any treatment as 100%. The residual rates of the final concentrations of inhibitory proteins in the three experimental groups, they were 30, 60 and 90 ug mL^−1^, respectively.

**TABLE 1.** The MICs and MBCs of fusion protein DN-AflR and DN

| | Concentrations ($\mu\text{g ml}^{-1}$) | | | | | | | MIC | MBC |
| --- | --- | --- | --- | --- | --- | --- | --- | --- | --- |
| | 30 | 50 | 100 | 150 | 400 | 600 | 900 | ($\mu\text{g/ml}$) | ($\mu\text{g/ml}$) |
| DN-AflR | + | + | + | - | - | - | - | 150 | 400 |
| DN | + | + | + | + | + | - | - | 600 | >900 |
| Negative control | + | + | + | + | + | + | + | / | / |

Negative control: no inhibitory protein.
+: observed fungal growth.
-: no fungal growth observed.
/ : no inhibition.

### Inhibitory effect of fusion protein on aflatoxin B_1_ biosynthesis

Two groups of experiments were performed using DN-AflR and DN with final concentrations of 30, 60, and 90 μg ml^−1^. The residual rate of aflatoxin B_1_ after fusion protein treatment was 24.37%, 14.21% and 6.74%, respectively, and the residual rate of aflatoxin B_1_ after DN treatment was 44.39%, 22.19% and 10.99%, respectively (Fig. 4B). Compared with DN, fusion protein had stronger inhibitory effect on biosynthesis of aflatoxin B_1_ (*P*<0.05). At 30 μg ml^−1^ of DN-AflR, more than 75% aflatoxin B_1_ was degraded.

### Physical properties and molecular structure of DN-AflR

Amino acid sequence, isoelectric point and other basic informations were analyzed by ExPASy (http://web.expasy.org/protparam/; Table 2). Secondary structure of protein was predicted by SPLIT (http://splitbioinf.pmfst.hr/split/4/; Fig. 5A). Spatial structure was predict by SWISS-MODEL. The fusion protein DN-AflR carries a positive charge, has a relative strong hydrophobic effect, and contains the characteristic secondary structure of the protein. By analyzing the predicted structures of DN-AflR (Fig. 5B) and GAL4 (Fig. 5C), DN-AflR has the zinc finger DNA-binding functional structure (specifically binds to AflR binding site) (Fig. 5D).

**FIG 5.**
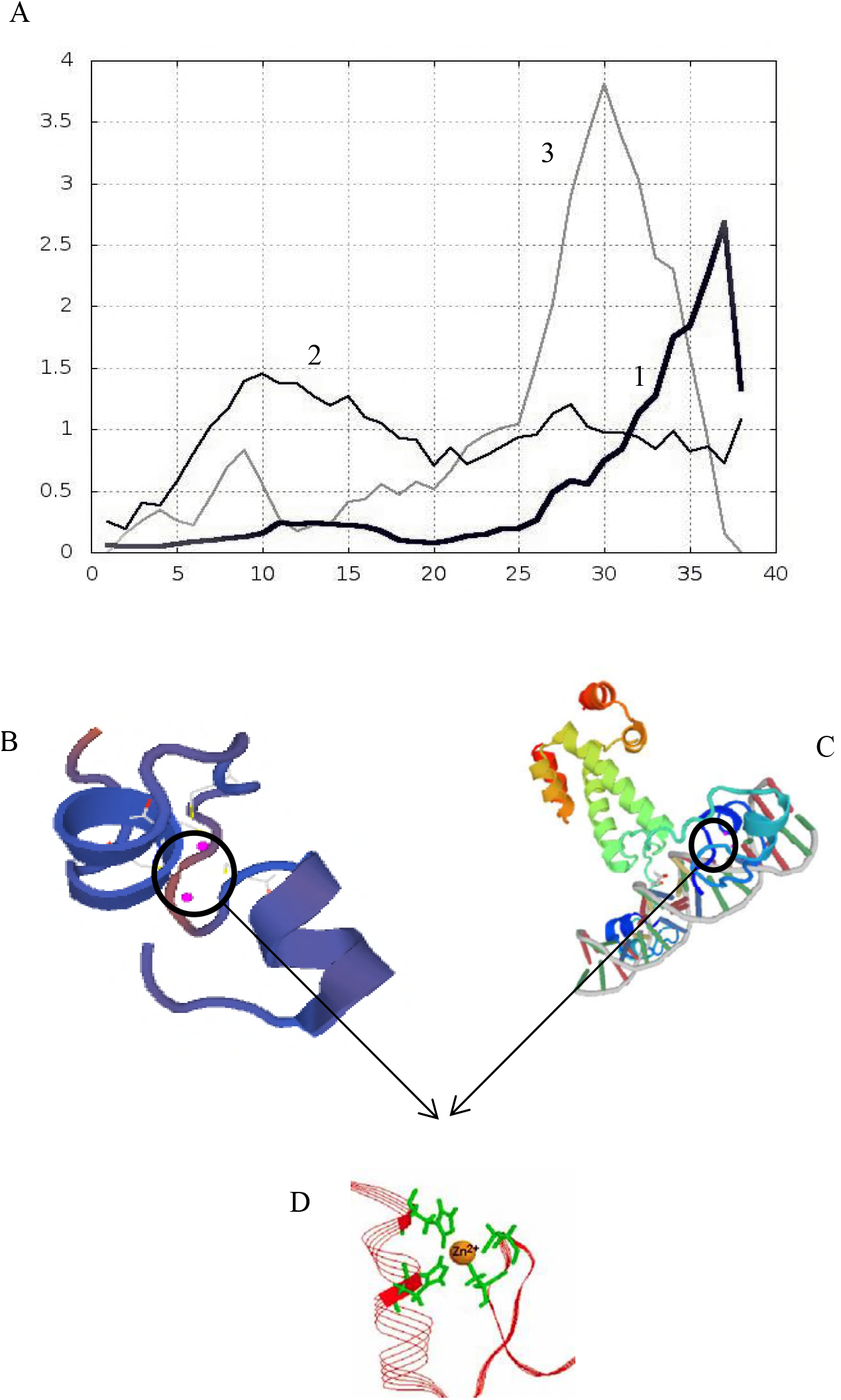
Predicted structure of DN-AflR. (A) Secondary structure. Line 1: Transmembrane helix preference (THM index). Line 2: Beta preference (BET index). Line 3: Modified hydrophobic moment index (INDA index). (B) Spatial structure of DN-AflR. Marked position in circle: zinc finger structure. (C) Structure of GAL4. Marked position in circle: zinc finger structure. (D) Zinc finger structure.

**TABLE 2.** The basic properties of DN-AflR

|  |  |
| --- | --- |
| Number of amino acids | 36 |
| Theoretical pI | 8.33 |
| Formula | C<sub>146</sub>H<sub>250</sub>N<sub>50</sub>O<sub>54</sub>S<sub>6</sub> |
| Total number of atoms: | 506 |
| Aliphatic index | 40.83 |
| Total number of negatively charged residues (Asp+Glu) | 4 |
| Total number of positively charged residues (Arg+Lys) | 6 |
| Grand average of hydropathicity (GRAVY) | -0.581 |

Basic informations were analyzed by ExPASy (http://web.expasy.org/protparam/).

## DISCUSSION

At present, more and more microorganisms and their metabolites were reported to inhibit the growth of *Aspergillus flavus* and degrade aflatoxins (5, 26). Albert et al. transformed the laccase gene of *T. versicolor* into recombinant *A. niger* by gene cloning, and the inhibition rate on aflatoxin of laccase (118 U L^−1^) after recombinant expression as high as 55% (27). Douillard et al. demonstrate that novel *L. lactis* fusion partner expression vectors allow high-level expression of soluble heterologous proteins (28). Study showed that Fh8 tag fusion expression could significantly improve the ability of *E. coli* to express soluble exogenous proteins (29). Genetic engineering could highly express the active proteins and other metabolites in prokaryotic or eukaryotic hosts, which is the great way to cut down costs.

However, there was few report on the inhibition of growth of *A. flavus* by molecular modification of positive regulatory genes in aflatoxin biosynthesis. Fusion expression has been continuously applied to the process of expressing recombinant protein in order to improve functions of the active protein (30, 31). TEM and SEM are effective applications to analyze the characteristics and morphologies of samples. SEM showed that DN caused changes in mycelial morphology of *A. flavus*. TEM showed the ruptured vacuoles accounted for most of cell space and destroyed the function and balance of other organelles, which affected normal growth of the whole cell (Fig. 2). So that DN had the ability of inhibiting the growth of *A. flavus*.

In order to improve the anti-*A. flavus* effect of DN, we fused the genes encoding DN and sequences of zinc finger cluster protein (specifically binds to the AflR binding site), then successfully constructed a recombinant plasmid dn-aflR-pET32a (Fig. 1). In the process of *E. coli* expression, when expressed in the host system at a high level, the recombinant protein was easy to form inclusion bodies (32). This may be due to the fact that during the expression, the protein folded too fast, while insufficient supply of enzymes or co-factors made it cannot form the correct secondary bond. According to the reports (33), we used 16°C, 160 rpm as the conditions for inducing expression. In general, target proteins with a molecular weight of less than 5 kDa or more than 100 kDa could not be expressed easily. The smaller molecular weight of protein, the easier it was to be degraded. In order to avoid degradation of target protein due to its low molecular weight, we used the pET32a vector to express the active protein, and its Trx and His tag significantly improved the stability of expression, and enhanced purification of protein, respectively. Because target protein contained the His tag, result of SDS-PAGE showed that molecular weight of fusion protein was 24 kDa (Fig 3), which was a little more than the predicted molecular weight by the theoretical calculations (20 kDa). Compared with DN, the fusion protein DN-AflR had stronger inhibitory effect on growth of *A. flavus* (Fig. 4A). Simultaneously, it also had significant advantages in inhibiting aflatoxin B_1_ biosynthesis (Fig. 4B), especially, under 30 μg mL^−1^ concentration of DN-AflR, more than 75% aflatoxin B_1_ was inhibited. To get a deeper comprehension of DN-AflR, its physical properties and molecular structures were predicted. Outer membrane of most cells is negatively charged, while most anti-fungal proteins are positively charged and they can bind to the cells surface through electrostatic attraction (34). The α-helical peptides with higher hydrophobic properties have a higher ability to lyse membranes, and the inhibitory activity of some peptides would even disappear with reduced hydrophobicity (35). Through the predicted results, the fusion protein DN-AflR carries positive charge and has a relative strong hydrophobic effect, and contains the secondary structure of helix. Especially, DN-AflR contains the zinc finger cluster protein structure (compare with GAL4), which make it could compete with AflS-AflR complex by acting on AflR binding sites to inhibit the activation of aflatoxin. These properties confirmed that DN-AflR had high inhibitory ability on growth of *A. flavus* and biosynthesis of aflatoxin.

To the best of our knowledge, this is the first report about fusing the genes encoding inhibitory peptides and specific zinc finger cluster protein, and fusion expressed the novel anti-*A. flavus* protein. Results showed the modified novel protein reduced aflatoxin biosynthesis and growth of *A. flavus*, which could control aflatoxin contamination. Compared with DN, the inhibitory ability of novel protein has been improved significantly.

This work promoted deep researches for fusion expression of anti-fungal proteins. Furthermore, this study also provided more theoretical basis and technical support for the further biocontrol of *A. flavus* and a new idea about enhancing the anti-*A. flavus* ability of inhibitory substances.

## MATERIALS AND METHODS

### Materials, strains, and culture conditions

*A. flavus* NRRL3357, preserved in School of Food Science and Engineering, Ocean University of China, was maintained at 4°C on potato dextrose agar (PDA; Bio-way technology, Shanghai, China). For liquid culture, *A. flavus* was transferred into a 250 ml Erlenmeyer flask containing 100 ml of MM medium at 28°C. *Escherichia coli* DH5α and *E. coli* BL21 (Ruibio Biotech, Beijing, China) were used as hosts for plasmid amplification and genes expression, respectively. *E. coli* was grown in LB medium (10 g l^−1^ Tryoton, 5 g l^−1^ Yeast Extract, 10 g l^−1^ NaCl) containing 50 μl ml^−1^ ampicillin (Solarbio, Beijing, China) at 37°C with shaking at 180 rpm.

### Effects of DN on mycelia morphology of *A. flavus*. (i) Preparation of spores suspension

Spores of *A. flavus* were washed off with sterile distilled water containing 0.1% Tween-80 from PDA medium and filtered with a cotton slag to prepare spores suspension. The number of spores was counted by haemacytometer and diluted to the desired concentration.

### (ii) TEM and SEM

0.8 mg ml^−1^ DN (synthesized in GL Biochem, Shanghai, China) and 10^5^ spores ml^−1^ were added in 50 ml MM medium and cultivated at 28°C with 180 rpm for 48 h. Control group didn’t contain DN. Then the mycelia were fixed with 2.5% glutaraldehyde, dehydrated in ethanol and embedded in an epoxy resin to be observed by Transmission Electron Microscope (TEM; JEOL, Tokyo, Japan) (21), and the mycelia were also observed with Scanning Electron Microscope (SEM; JEOL, Tokyo, Japan) after dispersion by ultrasonic wave (22).

### Expression and identification of fusion protein. (i) Expression vector and its construction and transformation into *E. coli* BL21

The fusion gene *dn-aflR* containing genes encoding DN and zinc binuclear cluster protein (Cys-Xaa2-Cys-Xaa6-Cys-Xaa6-Cys-Xaa2-Cys-Xaa6-Cys, based on AAM03003.1, NCBI) was synthesized by Hongxun Biotech (Suzhou, China), and amplified with PCR (forward primer 5’-GAATTCGACAACGACAACT-3’, reverse primer 5’-AAGCTTACAGGCAAGACCA-3’). Sequences for restriction site EcoR I were sequentially incorporated at the 5’-end and Hind III restriction site was in order added to the 3’-end, and extracted gene by Plasmid Mini Kit (OMEGA, Omega bio-tek, Shanghai, China). Restriction enzymes Hind III and EcoR I were used to digest the *dn-aflR* and pET32a (+), then the digested fragments were separated and identified on 1% agarose gel and recovered from gel using Agarose DNA Extraction Kit (OMEGA, Omega bio-tek, Shanghai, China). The digested fragments were ligated at 4°C with T4 DNA ligase (Thermo Fisher Scientific, New York, USA) (Fig. 1). Then recombinant plasmid dn-aflR-pET32a was transformed into *E. coli* DH5α and single colony was picked for colony PCR. The recombinant plasmid extracted from suspension and empty plasmid were both transformed into *E. coli* BL21 by heat shock, and cultivated on solid LB medium containing 50 μg ml^−1^ ampicillin (23). After 16 h, single colony was inoculated into LB liquid medium and sent it to Ruibio Biotech (Beijing, China) for DNA sequencing.

### (ii) Protein expression and analysis

A single colony (containing dn-aflR-pET32a) was inoculated into 10 ml LB medium and cultured at 37°C. Then overnight culture was inoculated into fresh 300 ml LB media in 1:5 ratio. When optical density (OD600) reached 0.6, the final concentration of 0.8 mM IPTG was added to induce the expression. After 10 h induction at 16°C under 160 rpm, the cells were washed with buffer (20 mM Tris, 30 mM NaCl, 10 mM imidazole, pH=7.5) and then lysed by sonication (JY92-IIN, Ningbo Scientz Biotech, Ningbo, China; ultrasonic power 300 w, ultrasound work 4 s, stop 6 s, 60 times). Supernatant was collected and identified by SDS-PAGE.

### (iii) Protein purification

The presence of a His tag in the recombinant protein meant that the purification was performed with Ni^2+^-NTA affinity chromatography. Briefly, samples were passed through a column and extensively washed with the elution buffer (100 mM Tris base, 500 mM NaCl, and 200 mM Imidazole, pH=8). Purified fusion protein was demonstrated by SDS-PAGE and relative quantity was determined by Braford protein assay in wave length of 595 nm by standard protein BSA (bovine serum albumin) (Solarbio, Beijing, China).

### Effects of DN and DN-AflR on growth of *A. flavus*. (i) Anti-*A. flavus* activity

Fusion protein was evaluated by agar disk diffusion experiment. The diameter of inhibition zone indicated the anti-*A. flavus* activity. 0.1 ml spores suspension (10^5^ spores ml^−1^) was coated on PDA plate, and Oxford Cups filled with 100 μl DN (600 μg ml^−1^) and DN-AflR (600 μg ml^−1^) were put on it, respectively. The Oxford Cup on control plate was filled with 100 μl sterile water.

### (ii) MIC and MFC

A series of solutions containing DN-AflR or DN at the concentrations of 30, 50, 100, 150, 400, 600 and 900 μg ml^−1^ were prepared by serial dilution. 200 μl of above solutions was added into each well of 96-well plates containing PDA medium, respectively. Pure PDA without DN-AflR and DN was the control. Then 5 μl suspension (10^5^ spores ml^−1^) was added to each well and placed at 28°C for 48 h (24). By definition, the minimum concentration of compound that completely inhibited growth of *A. flavus* was the minimum inhibitory concentration (MIC) (non-visible *Aspergillus* hyphae); the culture was continued for 8 days to get the MFC, while the lowest fungicidal concentration (MFC) was the lowest concentration which no spore could germinate at.

### Effects of DN and DN-AflR on AFB_1_ biosynthesis

DN or DN-AflR (final concentrations of 30, 60 and 90 ug ml^−1^) were added into 30 ml MM medium, respectively, and 100 μl of suspension (10^5^ spores ml^−1^) was inoculated to each group and incubated at 28°C with 200 rpm. Each experiment was repeated 3 times. After culturing for 48 hours, samples were centrifuged at 8000 g for 20 min at room temperature. Five milliliters of supernatant were diluted with 5 ml ultrapure water. Next, 5 ml of the diluted sample was extracted in immune affinity columns (Huaan Magnech BioTech Co., Ltd., Beijing, China) and then eluted with 1 ml of methanol at a flow rate of 1 drop per second. The eluent was evaporated under a gentle stream of nitrogen at 45°C up to dryness condition, and then derivatized with 200 L hexane and 100 L trifluoroacetic acid (TFA) for 15 min. After being evaporated to dryness again, the eluent was redissolved in 200 L water-acetonitrile (85:15, v/v).

AFB_1_ was analyzed according to retention time in HPLC system equipped with a ZORBAX Eclipse XDB-C18 column (4.6×150 mm, 5um, Agilent, Palo Alto, CA, USA) and a 470 fluorescent detector (G1321A, Agilent, USA) (λexc 360 nm; λem 440 nm) using a mobile phase solvent of 10% acetonitrile, 40% methanol, and 50% water. The flow rate was 0.8 ml min^−1^ and injection volume was 20 μl (25).

### Statistical analysis

All data were presented as mean ± S.D. One-way analysis of variance (ANOVA) and Duncan’s multiple comparison test were used and carried out with SPSS software (SPSS Inc., Chicago, IL, USA). *P*<0.05 was considered statistically significant.

## ACKNOWLEDGEMENTS

We thank National Natural Science Foundation of China (31471657) and the research grant from Qingdao Municipal Science and Technology Project (16-6-2-39-nsh) for supporting this research.

